# PEDIA: Prioritization of Exome Data by Image Analysis

**DOI:** 10.1101/473306

**Authors:** Tzung-Chien Hsieh, Martin Atta Mensah, Jean Tori Pantel, Krawitz Peter, the PEDIA consortium, Dione Aguilar, Omri Bar, Allan Bayat, Luis Becerra-Solano, Heidi Beate Bentzen, Saskia Biskup, Oleg Borisov, Oivind Braaten, Claudia Ciaccio, Marie Coutelier, Kirsten Cremer, Magdalena Danyel, Svenja Daschkey, Hilda David-Eden, Koenraad Devriendt, Sandra Dölken, Sofia Douzgou, Dejan Đukić, Nadja Ehmke, Christine Fauth, Björn Fischer-Zirnsak, Nicole Fleischer, Heinz Gabriel, Luitgard Graul-Neumann, Karen W. Gripp, Yaron Gurovich, Asya Gusina, Nechama Haddad, Nurulhuda Hajjir, Yair Hanani, Jakob Hertzberg, Hoertnagel Konstanze, Janelle Howell, Ivan Ivanovski, Angela Kaindl, Tom Kamphans, Susanne Kamphausen, Catherine Karimov, Hadil Kathom, Anna Keryan, Salma-Gamal Khalil, Alexej Knaus, Sebastian Köhler, Uwe Kornak, Alexander Lavrov, Maximilian Leitheiser, J. Gholson Lyon, Elisabeth Mangold, Purificación Marín Reina, Antonio Martinez Carrascal, Diana Mitter, Laura Morlan Herrador, Guy Nadav, Markus Nöthen, Alfredo Orrico, Claus-Eric Ott, Kristen Park, Borut Peterlin, Laura Pölsler, Annick Raas-Rothschild, Nicole Revencu, Christina Ringmann Fagerberg, Peter Nick Robinson, Stanislav Rosnev, Sabine Rudnik, Gorazd Rudolf, Ulrich Schatz, Anna Schossig, Max Schubach, Or Shanoon, Eamonn Sheridan, Pola Smirin-Yosef, Malte Spielmann, Eun-Kyung Suk, Yves Sznajer, Christian Thomas Thiel, Gundula Thiel, Alain Verloes, Irena Vrecar, Dagmar Wahl, Ingrid Weber, Korina Winter, Marzena Wiśniewska, Bernd Wollnik, Ming Wai Yeung, Max Zhao, Na Zhu, Johannes Zschocke, Stefan Mundlos, Denise Horn

**Affiliations:** Institute of Genomic Statistics and Bioinformatics, University of Bonn, Bonn, Germany; Charité – Universitätsmedizin Berlin, corporate member of Freie Universität Berlin, Humboldt-Universität zu Berlin, and Berlin Institute of Health, Institute of Medical Genetics and Human Genetics, Berlin, Germany; Monterrey Institute of Technology and Higher Education, Mexico; FDNA Inc., Boston Massachusetts, United States; Rigshospitalet, Department of Neurology, Copenhagen, Denmark; Unidad de Investigación Médica en Medicina Reproductiva, Mexico; University of Oslo, Oslo, Norway; CeGaT GmbH, Tübingen, Germany; University of Milan, Milan, Italy; Department of Human Genetics, University Hospital of Bonn, Bonn, Germany; Heinrich Heine University Düsseldorf, Düsseldorf, Germany; Catholic University Leuven, Leuven, Belgium; University of Hamburg, Hamburg, Germany; University of Manchester, Manchester, United Kingdom; Division of Human Genetics, Medical University of Innsbruck, Innsbruck, Austria; University of Tübingen, Tübingen, Germany; A. I. duPont Hospital for Children, Wilmington, United States; National Research and Applied Medicine Centre “Mother and Child”, Belarus; Lineagen, United States; Santa Maria Nuova Hospital, Italy; Center for Chronically Sick Children (Sozialpädiatrisches Zentrum, SPZ), Charité - Universitätsmedizin Berlin, Berlin, Germany; GeneTalk, Bonn, Germany; University Hospital Magdeburg, Magdeburg, Germany; Children’s Hospital of Los Angeles, Los Angeles, United States; Medical University of Sofia, Sofia, Bulgaria; Charité – Universitätsmedizin Berlin, corporate member of Freie Universität Berlin, Humboldt-Universität zu Berlin, and Berlin Institute of Health, NeuroCure Clinical Research Center, Berlin, Germany; Research Institute of Medical Genetics of Russian Academy of Medical Sciences, Russian Federation; Cold Spring Harbor Laboratory, Woodbury, United States; University of Bonn, Bonn, Germany; Hospital General Universitario De Valencia, Valencia, Spain; Hospital General De Requena, Servicio Pediatría, Spain; University Hospital Leipzig, Leipzig, Germany; Hospital Universitario Miguel Servet, Spain; Azienda Ospedaliera Universitaria Senese, Siena, Italy; Children’s Hospital Colorado, United States; Clinical Institute of Medical Genetics, University Medical Centre Ljubljana, Ljubljana, Slovenia; Sheba Medical Center, Israel; Université Catholique de Louvain, Bruxelles, Belgium; Odense University Hospital, Odense, Denmark; The Jackson Laboratory for Genomic Medicine, Farmington, United States; Berlin Institute of Health (BIH), Anna-Louisa-Karsch 2, 10178 Berlin, Germany; School of Medicine, University of Leeds, Leeds, United Kingdom; Ariel University, Ariel, Israel; Department of Genome Sciences, University of Washington, Seattle, United States; Center for Prenatal Diagnosis and Human Genetics, Berlin, Germany; Cliniques universitaires Saint Luc UCL, Bruxelles, Belgium; Institute of Human Genetics, Friedrich-Alexander-Universität Erlangen-Nürnberg FAU, Erlangen, Erlangen, Germany; Hopital Robert Debré, Paris, France; Center for Human Genetics and Laboratory Diagnostics, Germany; Poznañ University of Medical Sciences, Poznañ, Poland; University Medical Center Göttingen, Göttingen, Germany

## Abstract

Phenotype information is crucial for the interpretation of genomic variants. So far it has only been accessible for bioinformatics workflows after encoding into clinical terms by expert dysmorphologists. Here, we introduce an approach, driven by artificial intelligence that uses portrait photographs for the interpretation of clinical exome data. We measured the value added by computer-assisted image analysis to the diagnostic yield on a cohort consisting of 679 individuals with 105 different monogenic disorders. For each case in the cohort we compiled frontal photos, clinical features and the disease-causing mutations and simulated multiple exomes of different ethnic backgrounds. With the additional use of similarity scores from computer-assisted analysis of frontal photos, we were able to achieve a top-10-accuracy rate for the disease-causing gene of 99 %. As this performance is significantly higher than without the information from facial pattern recognition, we make gestalt scores available for prioritization via an API.

---

Rare diseases affect approximately 6% of the population, with genetic syndromes accounting for about 80 %.^1,2^The more than 5,000 entities represent a heterogeneous group of diseases, differing in cause, symptoms, and treatment, making diagnosis an important yet challenging healthcare issue. Due to extensive clinical variability this is true even for well characterized syndromes.^1,3^

Worldwide, more than half a million children born per year have a rare genetic disorder that is suitable for a diagnostic workup by exome sequencing, which has an unprecedented diagnostic yield for many indications such as developmental delay.^4–9^ The main remaining concern for the integration of exome sequencing into clinical routine is to increase the efficiency of genetic variant interpretation. Making phenotypic information – the observable, clinical presentation – computer-readable is key in solving this problem, and in providing clinicians with a much-needed tool for diagnosing genetic syndromes.^10^

To date, the most advanced exome prioritization algorithms combine deleteriousness scores for mutations with semantic similarity searches of the clinical description of a patient.^11–15^ The human phenotype ontology (HPO) with its extensive vocabulary has become the *lingua franca* for this purpose.^16^ However, semantic similarity searches presuppose that facial features can be named. A facial gestalt that is simply described in the literature as *typical* or *characteristic* of a certain disease is of little help for these approaches.

Beyond language, capturing indicative patterns by deep-learning approaches has recently gained attention in assessing facial dysmorphism.^17–21^Artificial neural networks are now able to quantify the similarities of patient photos to hundreds of disease entities and achieve accuracies that match or even surpass the level of dysmorphologists in certain tasks.^22–25^ For this reason tools such as Face2Gene are now used in addition to human expertise to guide the molecular testing and to interpret sequence variants. Here we investigate systematically whether facial image analysis can improve the evaluation of exome data and qualifies as a next-generation phenotyping technology for next-generation sequencing.^26^

## Results

We first present an overview about the approach to prioritize exome data by image analysis (PEDIA); a detailed description is provided in the Methods.

### PEDIA classifier

For the assessment of genetic variants, different sources of evidence have to be considered, from a populational, molecular, and phenotypic level. PEDIA is a Bayesian heuristic, that can be used to update the probability that a mutation in a gene is disease-causing, given the phenotypic information contained in a frontal photograph.

To build this classifier, we first measured the similarities of the facial gestalt to 216 specific diseases in 679 individuals with the convolutional neural network DeepGestalt.^21^ By this means, we were able to acquire scores for disorders with a single genetic etiology that quantify the PP4 criteria of the ACMG guidelines which is used for variant interpretation.^27,28^

In addition to DeepGestalt, we computed further prediction scores that are widely used on clinical features (Phenomizer, Boqa, Feature) and genetic variants (CADD) for all individuals of the PEDIA cohort (Supplemental Table 1). ^29,30,31^ With this data set we trained and tested a support vector machine that can be used to prioritize the genetic variants in a VCF files from exome sequencing.

### Gene prioritization

The term next-generation sequencing (NGS) implies the interrogation of all genes in a single assay. Similarly, the term next-generation phenotyping (NGP) refers to technology enabling similarity searches on a large set of disorders based on clinical patient records and medical imaging data. In order to increase the efficiency in diagnostics, we combined both approaches and benchmarked gene prioritization.

Similar to the performance readout in Gurovich et al., the identification of the disease-gene in exome data also represents a multiclass classification problem and the number of sequence variants in the coding part of the genome illustrates the complexity of the diagnostic assessment. In reference guided-resequencing, about 20,000-30,000 single nucleotide variants and small indels have to be considered. Although the majority of these variants can be removed as benign polymorphisms, rare and potentially disease-causing mutations in more than 100 genes remain in a typical case with a suspected monogenic disorder. When only a deleteriousness score such as CADD is used to rank these mutations, the disease-causing gene is in the top 10 in less than 46 % of the cases of the PEDIA cohort. This performance increases to a top-10-accuracy rate of up to 88 %, when semantic similarity scores are included that are based on HPO feature annotations. These prioritization approaches also represent the current state of the art in diagnostic laboratories for single exomes.^13,14^ The additional information contained in frontal photos of dysmorphic cases pushes the correct disease-gene to the top-10 in more than 99 % of the cases in the PEDIA cohort and in the DeepGestalt test set (Figure 1 B).

**Figure 1:**
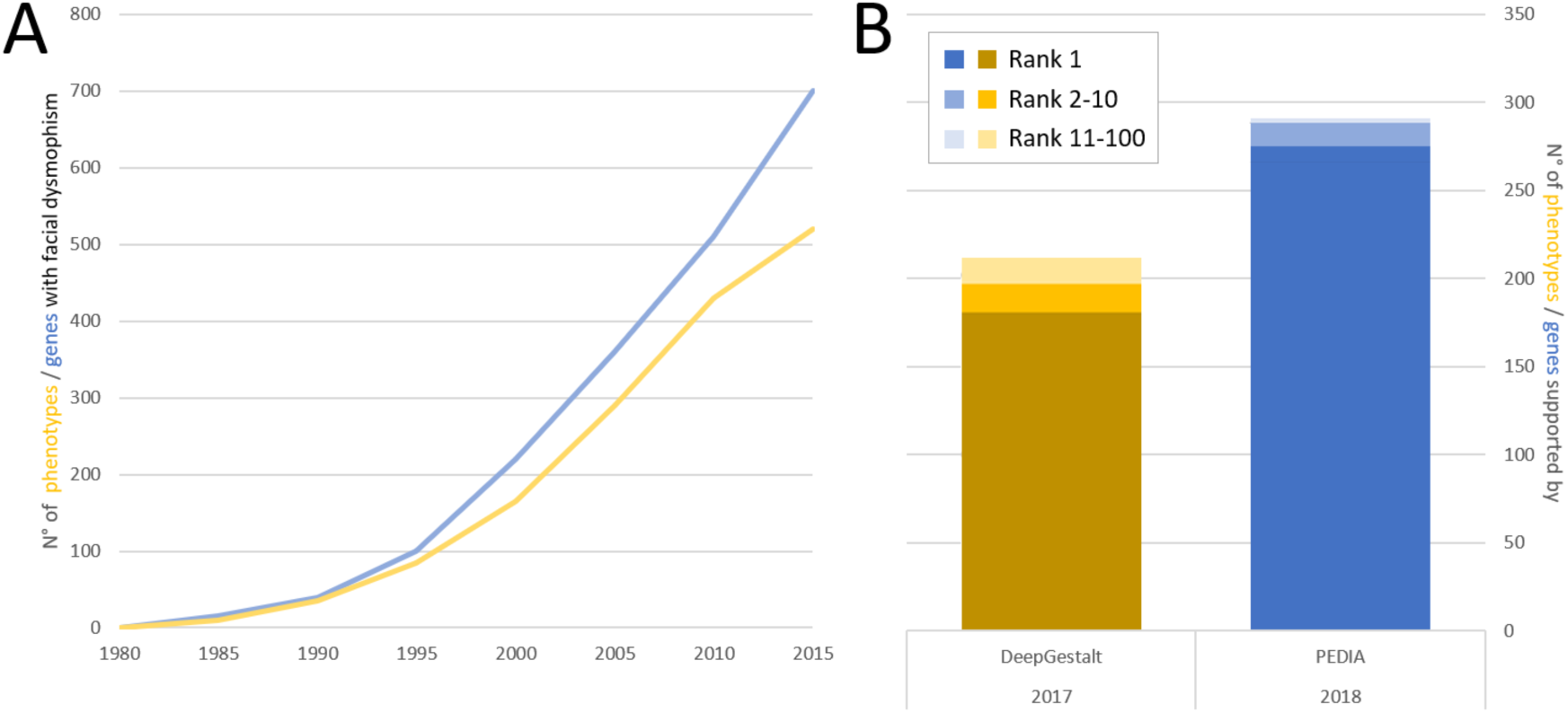
A) Schematic Increase of Mendelian phenotypes with facial abnormalities and associated genes listed in the encyclopedia of Online Mendelian Inheritance in Man over time. B) The next-generation phenotyping tool DeepGestalt could be used to differentiate between 216 disorders in the end of 2017 and achieved a top-10-accuracy rate of 90 %. The subset of Mendelian phenotypes that are suitable for a diagnostic workup by exome sequencing corresponds to 290 genes and in the PEDIA cohort a top-1-accuracy of 98 % was achieved.

The value of a frontal photograph can exemplarily be demonstrated by a case with Coffin-Siris syndrome that is shown in Figure 2 A: The characteristic facial features are relatively mild, so the correct diagnosis is only listed as the third suggestion by DeepGestalt. Amongst all the variants encountered in an exome data set, the disease-causing gene *ARID1B* would only achieve rank 24, if scored by the molecular information alone. However, in synopsis with the phenotypic information, the PEDIA approach lists this gene as first candidate by far (Figure 2 C).

**Figure 2:**
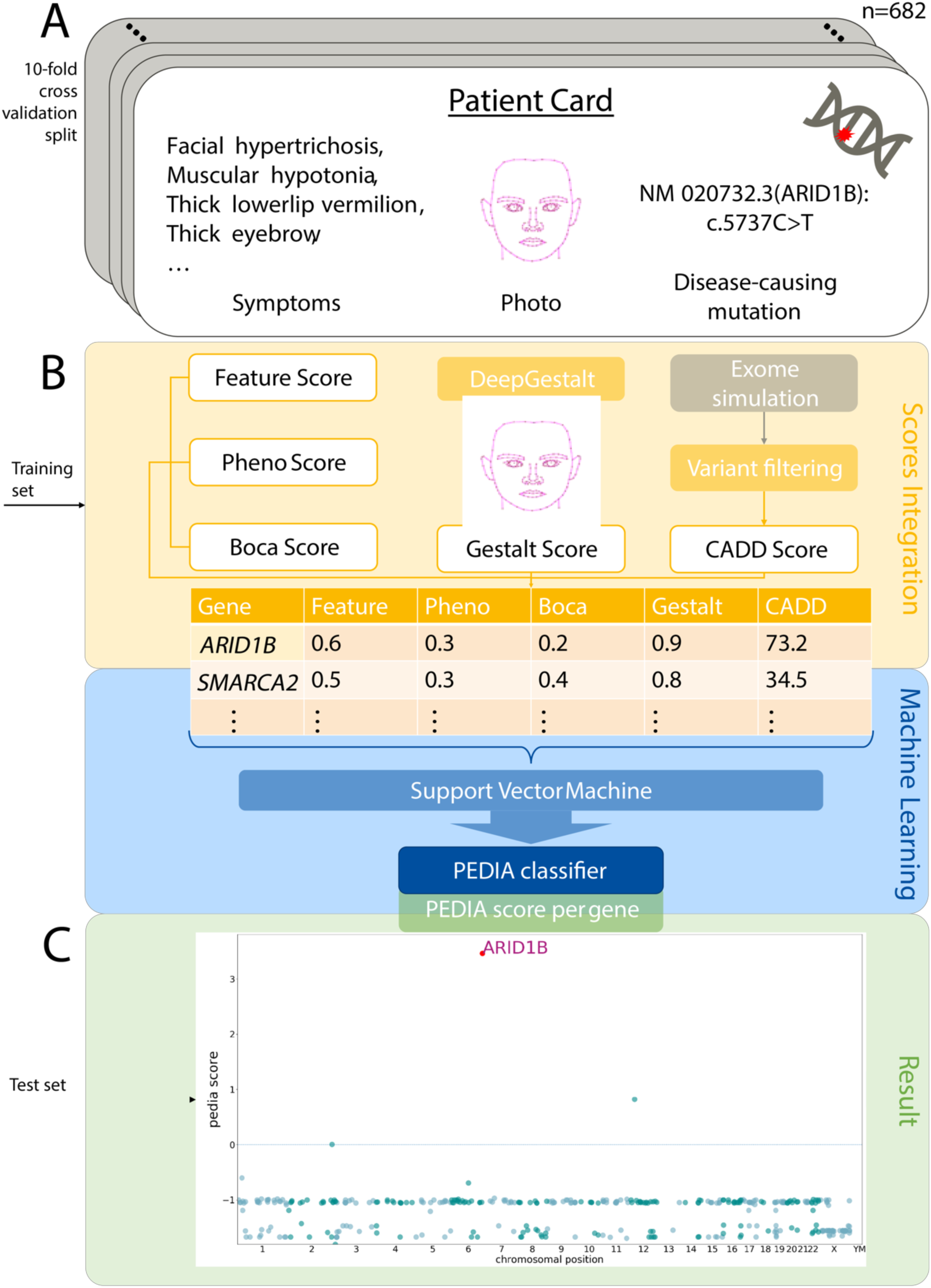
Prioritization of Exome Data by Image Analysis. A) Clinical features, facial photograph and disease-causing mutation of one individual of the PEDIA cohort. In total the cohort consists of 679 cases with monogenic disorders that are suitable for a diagnostic workup by exome sequencing. B) Clinical features, images and exome variants were evaluated separately and integrated to a single score by a machine learning approach. C) The disease-causing gene of the case depicted in A achieves the highest PEDIA score and molecularly confirms the diagnosis of Coffin-Siris syndrome. Other genes associated with similar phenotypes such as Nicolaides-Baraitser syndrome, achieved also scores for gestalt but not for variant deleteriousness. This figure has been adapted for bioRxiv by removing the patient photo. The original version with a patient photo is available on request. Also see https://pedia-study.org

Although the syndrome of the case shown in Figure 2 might also be molecularly confirmed by a directed single gene test in other instances where the facial gestalt is more indicative, the high phenotypic variability associated with disease-causing mutations is well-known for genes of syndromic disorders. It has been exhibited in the deciphering developmental disorders (DDD) project, that many such diagnoses were made only after exome sequencing.^6^ This finding is also reflected by frontal image analysis of the entire PEDIA cohort with DeepGestalt alone that achieves a top-10-accuracy rate for the disease-causing gene of around 58 %.

The efficiency of a prioritization algorithm can also be measured by the area under the curve (AUC) of the disease-causing mutation versus its ranked position. The higher the AUC, the higher the diagnostic yield in a fixed amount of time that is spend on the analysis of sequence variants (Figure 3). Combining similarity scores from image analysis, phenotypic features and molecular deleteriousness achieves the best AUC on the PEDIA cohort and is therefore suited to speed up diagnostics.

**Figure 3:**
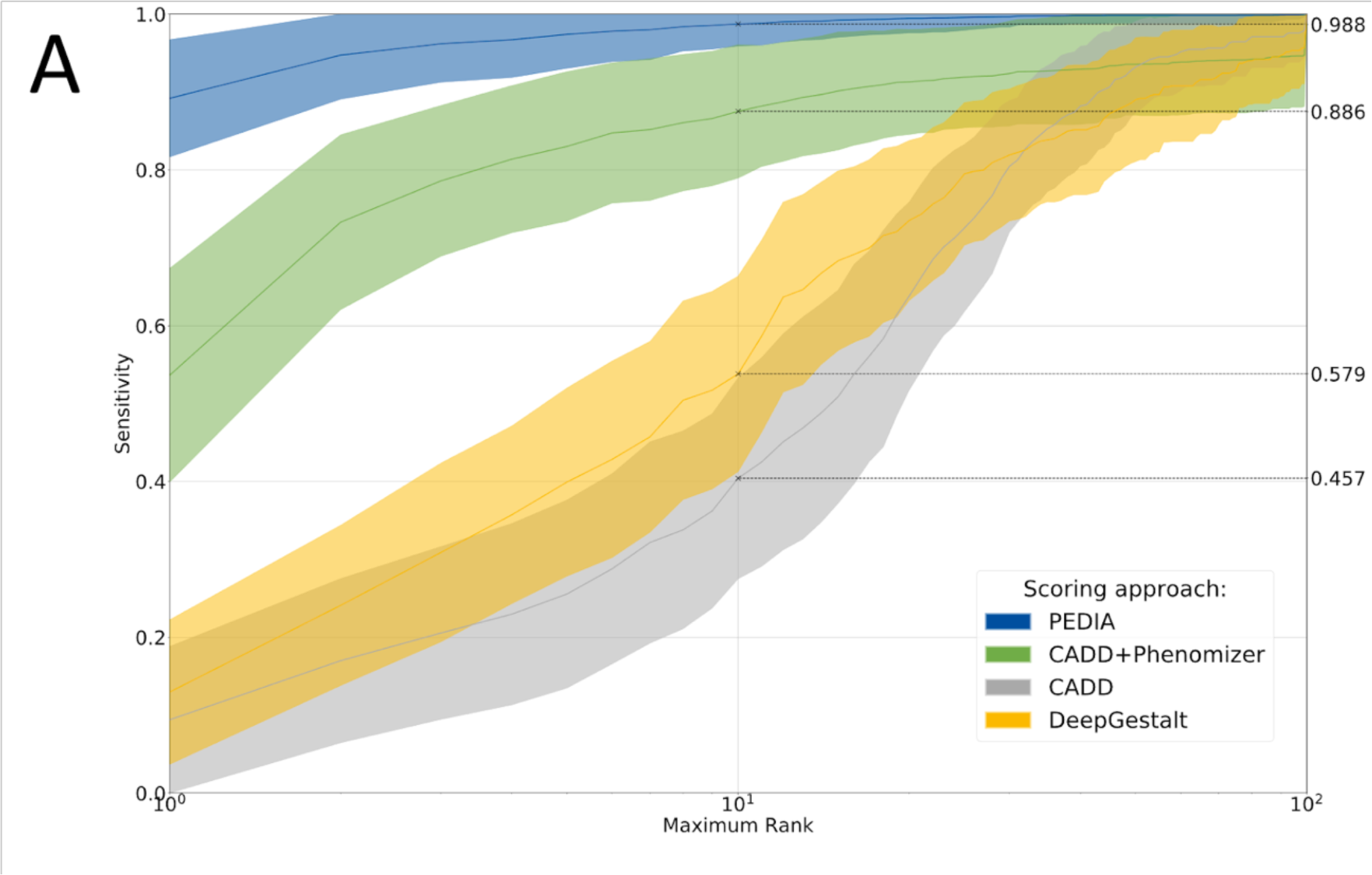
Area under the curve for different disease-gene prioritization approaches. For each case the exome variants are ordered according to four different scoring approaches, solely by a molecular deleteriousness score (C), by score from image analysis (DeepGestalt), by a combination of a molecular deleteriousness score and a clinical feature based semantic similarity score (P+C), or the PEDIA score that includes all three levels of evidence. The sensitivity of the prioritization approach depends on the number of genes that are considered in an ordered list. The top 10 accuracy rates of of Figure 1B correspond to the intersect of the curves for PEDIA and DeepGestalt at maximum rank 10^1^. Note that for benchmarking DeepGestalt on the gene level, syndrome similarity scores first have to be mapped to the gene level, resulting in a lower performance compared to the readout on a phenotype level, due to heterogeneity. The area under the curve is largest for PEDIA scoring. When e.g. the first ten candidate genes are considered, the syndromic similarity quantified by image analysis increases the sensitivity by about 20 % compared to P+C.

The contribution from the different sources of evidence to the PEDIA score is also reflected by the relative weight of the deleteriousness of the mutation (0.44), all feature-based scores combined (0.25) and the results from image analysis by DeepGestalt (0.31) that can be derived from a linear SVM model. We therefore also conclude that the information contained in a frontal photograph of patient goes beyond, what clinical terms can capture.

## Discussion

According to the current version of the Online Mendelian Inheritance of Man Catalog, mutations in about 4000 genes are linked to phenotypes that are often difficult to distinguish and diagnose by clinical features alone, making next-generation sequencing a key technology for their molecular confirmation. However, the size and high variability of the genome as well as the low prevalence of disease-causing variants – many of them occur *de novo* – explain why sequence data analysis of a single individual is still challenging and time consuming.^5,6^

The guidelines for variant classification in the laboratory follow a qualitative heuristic that combines distinct types of evidence (functional, population, phenotype, etc.) and is compatible with Bayesian statistics.^32^ The advantage of such a framework is that continuous evidence types can be integrated into the classification system. While *in silico* predictions about a variant’s pathogenicity have a relatively long history in bioinformatics and machine learning, the quantification of phenotypic raw data with systems of artificial intelligence just began. Analogous to a score for the deleteriousness of a gene variant, one can include the phenotypic similarity to a distinct syndrome caused by mutations in the respective gene.

We analyzed this approach in the PEDIA cohort, consisting of 679 cases and covering 105 distinct disorders mapping to 181 disease-genes. Among these disorders were 73 phenotypes for which the performance of facial image analysis alone has recently been evaluated.^21^ Although the top-10-accuracies for gestalt- and PEDIA-scoring cannot be compared directly, both approaches operate on a similar order of phenotypes and genes, respectively. Adding suitable molecular information to 260 cases from the DeepGestalt publication test set increased the correct disease-gene in the top 10 to about 99%, from 90% with only the phenotypic information. Considering only molecular information and clinical features, but without the results from image analysis, the correct disease gene would have only been placed in the top 10 in 62%. The genetic background, which might correspond to a different number of variant calls or higher load of deleterious mutations, had negligible influence on the performance.

The performance for the entire PEDIA cohort is comparable to the DeepGestalt test set. However, there are three important lessons learned from specific subgroups or cases achieving lower PEDIA ranks: 1) Although the convolutional neural network used for image analysis has been pretrained on real-world uncontrolled 2D images, patient photographs that were true frontal, of high resolution, with good lightening and contrast, and few artifacts such as glasses performed better. 2) Particularly rare diseases, or recently described disorders, for which the classifier’s representation is based on a smaller training set, show a lower performance, even if experienced dysmorphologists would consider them highly distinguishable.^24,34^ 3) Molecular pathway diseases, modeled as a single class, can be biased towards the prevailing gene if there is substructure in the phenotypic series, meaning there actually are gene-specific differences in the gestalt and complete heterogeneity is simply an approximation.^25^ This applies also to microdeletion syndromes that can be caused by single gene mutations, such as Smith-Magenis syndrome, or any clinical presentation of a phenotype that is considered atypical.

The only way to overcome the biases of semantic similarity metrics as well as AI-driven image analysis that are due to limited cohort sizes, is sharing of the phenotypic data sets.

In conclusion, the PEDIA study documents that exome variant interpretation benefits from computer-assisted image analysis of facial photographs, particularly if dysmorphism has been stated in the clinical notes. By including similarity scores from DeepGestalt, we improved the top-10-accuracy rate considerably. AI-driven pattern recognition of frontal facial patient photographs is an example of next-generation phenotyping technology with proven clinical value in the interpretation of next-generation sequencing data.

As deep-learning advances in the assessment of other medical imaging data, it will be interesting to study how these classifiers affect variant interpretation separately and in aggregate.^35,36^

## Supporting information

## Data and Code Availability

PEDIA is freely available for academic use at https://pedia-study.org and the source code is available at https://github.com/PEDIA-Charite.

## Acknowledgements

This work was funded by the Deutsche Forschungsgemeinschaft (KR 3985/7-3, KR 3985/6-1)

## Author contributions

Conceived and designed the study and drafted the manuscript: T.C.H., M.A.M., J.T.P., and P.M.K.

Project Coordination: M.A.M., J.T.P., N.F.,

Acquired, analyzed, and interpreted the clinical data: A.D., L.B.S., A.B., S.B., O.B., A.M.C., C.C., M.C, K.C., S.D., M.D., K.D., S.D., S.D., D.D., N.E., C.R.F., B.F.Z., H.G., K.G., Y.G., N.H., N.H., L.M.H., K.H., I.I.,J.H., A.K., C.K., H.K., S.K., A.K., A.K., U.K., A.L., M.L., G.L., E.M., D.M., A.O., K.P., B.P., L.P., P.M.R, N.R., S.R., A.R.R., S.R., G.R., U.S., A.S., P.S.Y., E.K.S., M.S., Y.S., C.T., G.T., A.V., I.V., D.W., I.W., K.W., M.W., B.W., M.W.Y., L.G.N., C.E.O.

Chief clinical data review: D.H.

Performed the Bioinformatics and statistical analysis: M.S., J.H., M.A.M., T.C.H., T.K., S.K., M.Z., N.Z., O.B., G.N., Y.G., Y.H., O.S., H.D.E., J.T.P., S.G.K.

Critically revised the manuscript for important intellectual content. H.B.B., P.N.R., S.M., J.Z.,

## Competing interests

N.F., H.D.E., Y.G., G.N., O.B., Y.H., are employees of FDNA Inc, T.K. is employee of GeneTalk GmbH.

## Materials and Methods

### Patients

We compiled a cohort of 679 patients with a Mendelian disorder to evaluate Prioritization of Exome Data by Image Analysis (PEDIA). For all cases in this cohort frontal facial photographs were available for analysis and clinical features were documented in HPO terminology.^16^ The diagnoses of all individuals have previously been confirmed molecularly and are suitable for analysis by exome sequencing. In total, the cohort covers 105 different monogenic syndromes that are linked to 181 different genes. Of the individuals in this cohort, 375 were published and 309 have not been previously reported (see Supplementary Appendix).

The study was approved by the ethics committees of the Charité - Universitätsmedizin Berlin and of the University of Bonn Medical Center. Written informed consent was provided by the patients or their guardians.

In addition to PEDIA data set, we analyzed a subset of the DeepGestalt study comprising all 260 cases of the publication set with monogenic syndromes which diagnosable by exome sequencing.^21^

### Data Preparation

The facial images were analyzed with DeepGestalt (FDNA), a deep convolutional neural network that was trained on more than 17,000 patient images.^21^ The results of this analysis are gestalt scores quantifying the similarity to 216 different rare phenotypes per individual. Although DeepGestalt is built as a framework that aims to learn from every additional case, we excluded all data of the PEDIA cohort from the model for benchmarking purposes in a similar manner as described in the original publication. In addition to the image analysis, we performed semantic similarity searches with the annotated HPO terms by Feature Match (FDNA), Phenomizer and BOQA.^29,30^

We filtered all sequence variants as described by Wright et al. and scored the remaining mutations for deleteriousness with CADD.^31,32^ If no exome data was available, we spiked the disease-causing mutation into the exome data of a healthy individual from the 1000 Genomes Project.^33^ This exome simulation was applied to the entire PEDIA cohort to assess the influence of the genetic background on the performance of our scoring approach.

For the variants remaining after filtering, we derived the similarity scores from image analysis and semantic similarity searches that were based on HPO feature annotations for the syndromes associated with the respective genes. If there were several syndromes linked to a single gene, the highest gestalt and feature scores were selected. Case data is represented as table with a variable number of lines representing genes and five columns for the different scores (Figure 1 B). All five scores with per line as well as the Boolean label disease gene “true” or “false” were used to train a classifier that yields a single value per gene, the PEDIA score, that can be used for prioritization (Figure 1 C). A detailed description of preprocessing and filtering, as well as all the annotated data, can be found in our code repository.

### Gene prioritization

We used a support vector machine (SVM) to prioritize the disease-causing gene in each patient. First, we split the PEDIA cohort into a training and a test set. We used a linear kernel on the five scores to train the SVM and selected the hyperparameter C in the range from 2^-6^ to 2^12^ by performing internal 5-fold cross-validation on the training set. The C with highest top-1 accuracy was selected for training linear SVM. We further benchmarked the performance of each case in the test set with this model. The distance of each gene to the hyperplane - defined as the PEDIA score - was used to rank the genes for the case. If the disease-causing gene was at the first position, we called it a top 1 match, or if it was amongst the first ten genes, we called it a top 10 match.

To evaluate the accuracy, we conducted a 10-fold cross-validation, that is, we split 679 cases into 10 groups to minimize overfitting. For the 260 cases from the DeepGestalt publication test set, where exome diagnostics would be applicable, we randomly selected a patient from the PEDIA cohort with the same diagnosis and replaced the entire gestalt scores per case. Thus, we were also able to analyze the influence of another large collection of patient images in the exome prioritization. In total, all experiments were conducted ten times and the achieved top-1 and top-10-accuracies were averaged. All training data as well as the classifier are available at https://github.com/PEDIA-Charite and https://pedia-study.org

### Performance evaluation in a classification task

Both, DeepGestalt and PEDIA are approaches to solve multiclass classification problems (MCPs), the first tool operating on phenotypes and the second on genes. The difficulty of the task is characterized by the number of classes and the distinguishability of the different entities. For both MCPs the maximum number of classes can be estimated from Online Mendelian Inheritance in Man catalog, that is currently listing around 500 distinct disorders with facial abnormalities and 700 corresponding genes with disease-causing mutations (Figure 1 A).

Learning a phenotype in a neural network requires a certain number of unrelated cases. By the end of 2017, DeepGestalt could distinguish between 216 different entities. Due to more training data, 60 new disorders were added in the last six months and the number is expected to increase further on.

The performance of a prioritization tool can be assessed by the proportion of cases in a test set for which the correct diagnosis or disease-gene is placed at the first position or amongst the first ten suggestions (top-1 and top-10-accuracy). The composition of the test set has an influence on the accuracy because some disease phenotypes are easier to recognize and some gene mutations are more readily identified as deleterious. The setup of the PEDIA cohort, which is comprehensively documented in the Supplementary Appendix, therefore aims at emulating the whole spectrum of cases that could currently be analyzed with DeepGestalt and diagnosed by exome sequencing.

